# Revisiting AXL as a therapeutic target in soft tissue sarcoma: transcriptomic profiling reveals largely AXL-independent activity of dubermatinib and bemcentinib

**DOI:** 10.64898/2026.10.02.755908

**Authors:** Cristina Garnés-García, Pablo Fernández-Aroca, Francisco J Cimas, María José Ruiz-Hidalgo, Borja Belandia, Noelia Mayor-Garnacho, Antonio Fernández-Aramburo, Victor Javier Sánchez-Arévalo Lobo, Ricardo Sánchez-Prieto, Diego M. Fernández-Aroca

**Affiliations:** Universidad de Castilla-La Mancha, Laboratorio de Oncología Molecular, Unidad de Medicina Molecular, Instituto de Biomedicina, Unidad Asociada de Biomedicina UCLM, Unidad Asociada al CSIC, Albacete, Spain; Instituto de Investigación Sanitaria de Castilla-La Mancha (IDISCAM).; Universidad de Castilla-La Mancha, Departamento de Química Inorgánica, Orgánica y Bioquímica, Área de Bioquímica y Biología Molecular. Facultad de Medicina, Albacete, Spain; Departamento de Biología del Cáncer, Instituto de Investigaciones Biomédicas “Sols-Morreale” (CSIC-UAM), Unidad Asociada de Biomedicina UCLM, Unidad Asociada al CSIC, Madrid, Spain; Translational Cancer Research Group, Chronic Diseases and Cancer, Area 3, Instituto Ramón y Cajal de Investigación Sanitaria (IRYCIS), Madrid, Spain; Sección de Oncología Médica, Complejo Hospitalario Universitario de Albacete, Albacete, Spain; Grupo de Oncología Molecular, Instituto de Investigaciones Biosanitarias, Facultad de Ciencias Experimentales, Universidad Francisco de Vitoria (UFV), Madrid, Spain; Grupo de Oncología Cutánea. Servicio de Anatomía Patológica, Hospital Universitario 12 de Octubre. Instituto de Investigación Sanitaria Hospital 12 de Octubre (imas12), Madrid, Spain

**Author notes:** Both authors have contributed by equal to this work (Cristina Garnés-García, Pablo Fernández-Aroca). Both senior authors have contributed by equal to this work (Ricardo Sánchez-Prieto, Diego M. Fernández-Aroca).

**Keywords:** 1. Keywords: Sarcoma, functional genomics, transcriptomics, AXL, kinase inhibitors, genetic knockdown

## Abstract

Soft tissue sarcomas (STS) are a heterogeneous group of mesenchymal malignancies with generally poor prognosis, raising a continued interest in novel therapeutic targets. The TAM-family receptor tyrosine kinase AXL has been proposed as a therapeutic target across several cancer types, including STS. Here we revisit the clinical relevance of AXL in STS by combining in silico analyses with functional and transcriptomic characterisation of two AXL inhibitors, dubermatinib and bemcentinib, in HT-1080 and SK-LMS-1 cells. STS ranked among the tumour types with the highest median AXL mRNA expression across The Cancer Genome Atlas, yet AXL expression was not associated with patient survival. Both inhibitors reduced cell viability and induced apoptosis but, unexpectedly, shRNA-mediated AXL knockdown didn’t reproduce their antiproliferative effect nor modified sensitivity to either drug. Transcriptomic profiling showed weak correlation between knockdown- and drug-induced changes, with most drug-regulated genes being AXL-independent and largely cell line-specific. Connectivity analysis against LINCS, regulatory-network inference and CRISPR dependency data instead point to mitotic and PI3K-mTOR pathways as candidate mediators of the therapeutic activity. Thus, dubermatinib and bemcentinib show antitumour activity in STS models that is largely independent of AXL inhibition, suggesting that the rationale for AXL-directed therapy may be reconsidered.

## 3. Introduction

Soft tissue sarcoma (STS) comprises a heterogeneous group of malignant tumours of mesenchymal origin, encompassing multiple histological subtypes with different molecular and clinical features. Despite advances in cancer therapy, the prognosis of patients with advanced or metastatic STS disease remains poor, underscoring the need for novel therapeutic targets and predictive biomarkers [1]. A growing body of evidence indicates that the TAM family of receptor tyrosine kinases (TYRO3, AXL and MERTK) has emerged as a key regulator of tumour progression, therapeutic resistance and immune evasion, with corresponding therapeutic potential [2]. Among the three TAM receptors, AXL has attracted particular attention because of its involvement in cell proliferation, migration, invasion, epithelial-to-mesenchymal transition and drug resistance across multiple tumour types [3].

A link between AXL and sarcoma biology has been established from several perspectives. *AXL* is highly expressed and has been proposed as a candidate therapeutic target in dedifferentiated and pleomorphic liposarcoma [4], as well as in leiomyosarcoma, where *TYRO3* and *AXL* expression supports their functional involvement and pharmacological potential [5]. Activating mutations in *AXL*, such as the recently described W451C variant, have also been identified in a subtype-specific manner in myxofibrosarcoma, providing direct genetic evidence of the contribution of AXL to oncogenesis in this disease [6]. Beyond its role as a biomarker, AXL may also have a role in the therapeutic response in sarcoma as, for instance, in Ewing sarcoma, where chemotherapy induces AXL activation and the combined inhibition of the JAK1/STAT6/GAS6/TAM axis improves chemotherapy efficacy [7], in agreement with previous works showing antiproliferative activity of the selective AXL inhibitor BGB324 (bemcentinib) [8]. Multi-kinase inhibitors with anti-AXL activity have similarly shown preclinical potential in Ewing sarcoma [9]. In parallel, proteomic profiling identified AXL activation as a candidate mechanism of acquired resistance to pazopanib in synovial sarcoma, reinforcing the rationale for combined or sequential AXL inhibition in cancer therapy [10].

Pharmacological interest in AXL has translated into the development of several small-molecule inhibitors. Bemcentinib (R428/BGB324) was among the first selective AXL inhibitors characterised, showing potent antimetastatic activity and extended survival in animal models of metastatic breast cancer [11], and has since progressed into early-phase clinical evaluation, both as monotherapy and in combination with other agents. More recently, dubermatinib (TP-0903) [12] has emerged as a second-generation AXL/GAS6-axis inhibitor that interferes with PI3K/AKT and MAPK/ERK signalling, promoting cell-cycle arrest and apoptosis while reducing proliferation, invasion and therapy resistance in several preclinical tumour models, including pancreatic cancer [13] and neuroblastoma [14].

In this context, we aimed to analyse the role of dubermatinib and bemcentinib in experimental models of STS at both the biological and transcriptomic level, using HT-1080 (fibrosarcoma) and SK-LMS-1 (leiomyosarcoma) cell lines. Our results confirm that both inhibitors show cytotoxic and pro-apoptotic effects in STS cells. However, shRNA-mediated *AXL* knockdown did not recapitulate the effects of either inhibitor on cell growth or viability, nor modified cellular sensitivity to either drug. A comparison of the transcriptional programmes induced by genetic *AXL* depletion and by pharmacological AXL inhibition revealed only limited overlap between drug- and knockdown-induced transcriptional changes, with the majority of drug-regulated genes being AXL-independent. In summary, although our data support a potential antitumour use for dubermatinib and bemcentinib in STS, they also indicate that AXL itself should be reconsidered as the principal driver of this activity, and raise the question of which additional targets mediate the observed effects.

## 4. Results

### 4.1.1. AXL is highly expressed and frequently amplified, but not prognostic, in STS

We first examined the status of *AXL* in STS using pan-cancer clinical and genomic data from The Cancer Genome Atlas (TCGA). STS ranked among the tumour types with the highest median *AXL* mRNA expression (Fig. 1A) and was among the TCGA cohorts with the highest overall frequency of *AXL* genetic alterations, driven predominantly by copy-number amplification (Fig. 1B). In contrast, *AXL* point mutations were rare (<2% of cases) and no *AXL* structural variants were detected in this cohort. Among STS histological subtypes, undifferentiated pleomorphic sarcoma (UPS) and myxofibrosarcoma showed the highest overall burden of AXL genetic alterations (Fig. S1).

**Fig. 1.**
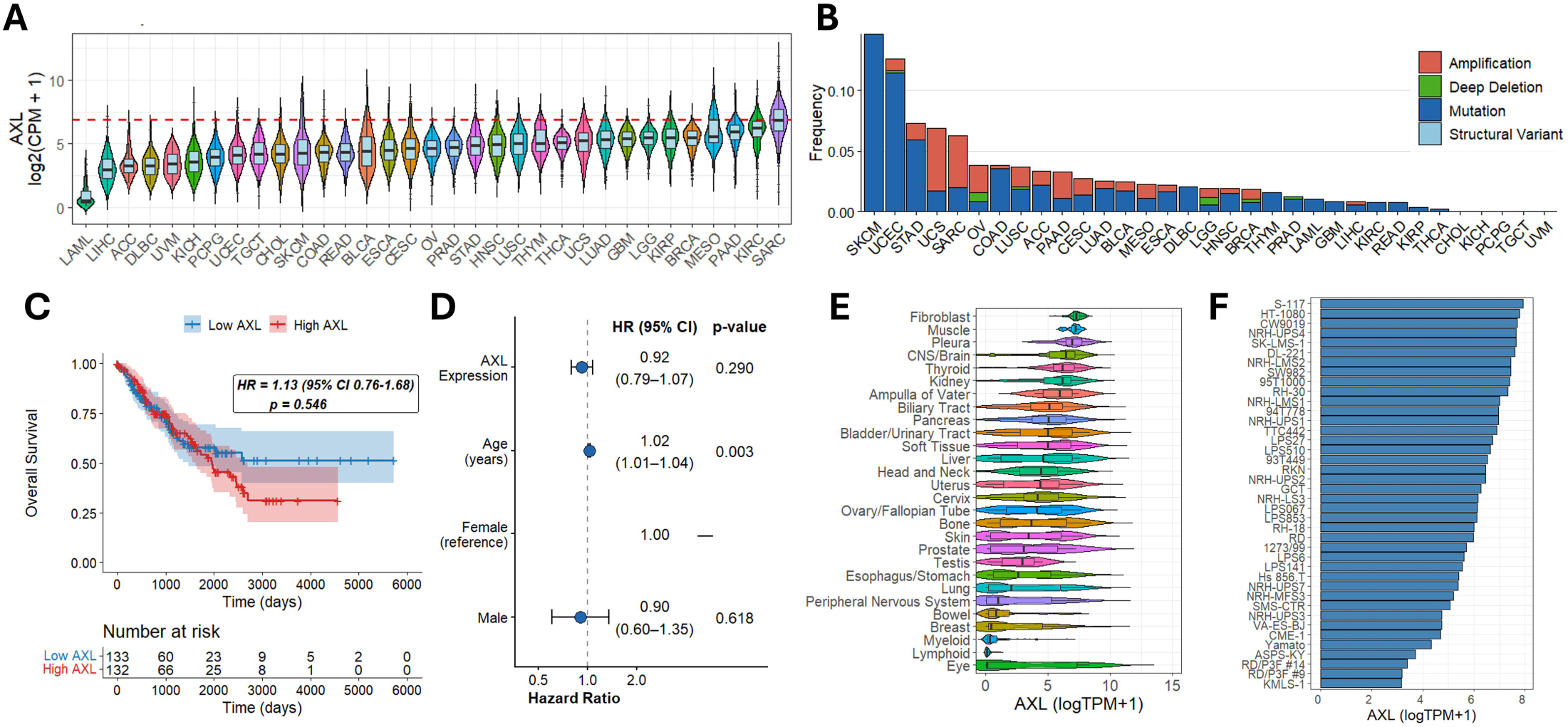
AXL expression, genetic alteration and prognostic value across TCGA tumour types and in STS. A) AXL mRNA median expression (log2[CPM+1]) across TCGA tumour types. Red dashed line indicates the TCGA-SARC median. B) Frequency of AXL genetic alterations across TCGA tumour types. C) Kaplan-Meier overall survival curves for TCGA-SARC patients stratified by median AXL expression. D) Forest plot of multivariable Cox model of overall survival including AXL expression, age and sex. E) Violin plot of AXL expression (log[TPM+1]) across CCLE cell line lineages. F) AXL expression across STS-lineage cell lines in CCLE.

Then, we asked whether *AXL* expression has prognostic value in STS by stratifying TCGA-SARC patients depending on the median levels of AXL. High AXL expression was not associated with overall survival (HR = 1.13, 95% CI 0.76–1.68, p = 0.546; Fig. 1C), which we confirmed with a multivariable Cox model including age, sex and *AXL* gene expression as a continuous variable (HR = 0.92, 95% CI 0.79–1.07, p = 0.290; Fig. 1D). Not surprisingly, age showed a subtle but significant association with survival (HR = 1.02, p = 0.003). This suggests that, despite its high expression in STS, AXL may not be considered a valuable prognostic biomarker in this tumour type.

To make an informed selection of the optimal cell lines for subsequent *in vitro* functional assays, we next evaluated *AXL* expression across cell lines from different lineages in the Cancer Cell Line Encyclopedia (CCLE), which revealed a wide range of *AXL* levels across the STS lineage (Fig. 1E). Within the STS lineage specifically, HT-1080 (fibrosarcoma) and SK-LMS-1 (leiomyosarcoma) were among the commercially available cell lines with the highest *AXL* expression (Fig. 1F), and were selected as representative models for functional characterisation of AXL-targeted therapy.

### 4.1.2. Dubermatinib and bemcentinib reduce viability and induce apoptosis in STS cell lines

We then evaluated the activity of two representative small-molecule AXL inhibitors, dubermatinib and bemcentinib, in HT-1080 and SK-LMS-1 cell lines expressing AXL protein levels to a similar extent (Fig. 2A). Dose-response assays revealed that dubermatinib reduced cell viability in both models in the nanomolar range, with SK-LMS-1 showing a slightly more resistant phenotype than HT-1080 (Fig. 2B). Bemcentinib showed a similar trend, but its activity was observed at the micromolar range (Fig. 2C), pointing to a marked difference in potency between the two compounds despite their common putative target. Importantly, the relative resistance of SK-LMS-1 to both drugs could not be explained by differences in AXL expression levels between the two cell lines. Similarly, dubermatinib and bemcentinib induced apoptosis in both models, albeit to a lesser level in SK-LMS-1 (Fig. 2D, Fig. S2), consistent with a degree of cell line-specific variations in the response to drug treatments. In contrast, both inhibitors reduced proliferation in each cell line to a similar extent (Fig. 2E-F).

**Fig. 2.**
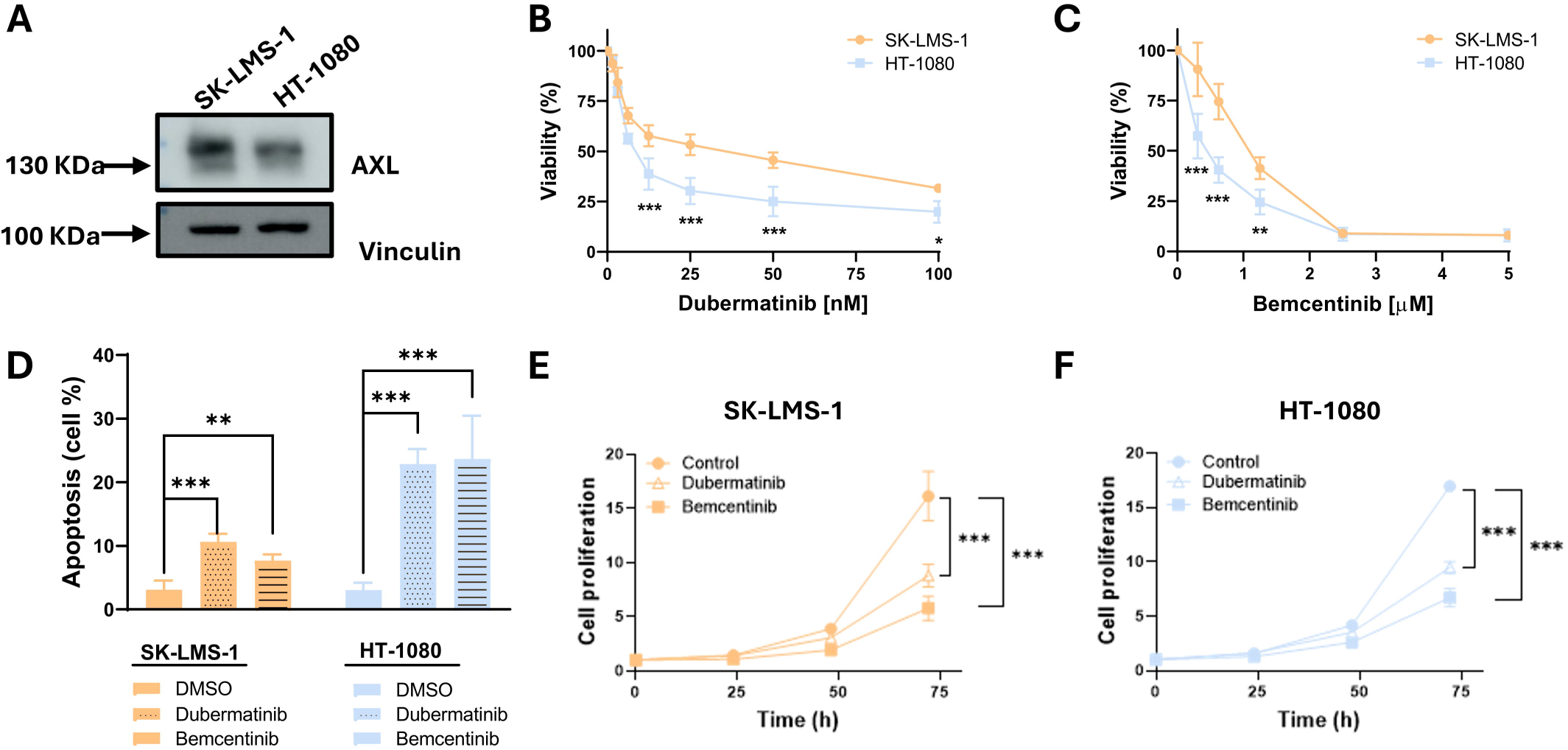
Dubermatinib and bemcentinib reduce viability and induce apoptosis in HT-1080 and SK-LMS-1 cells. A) AXL and vinculin (loading control) western blot in SK-LMS-1 and HT-1080 cells. B) Dose-response viability curves of HT-1080 and SK-LMS-1 cells treated for 72 h with indicated doses of dubermatinib. C) Same as in B for bemcentinib. D) Apoptosis induction quantification in SK-LMS-1 and HT-1080 cells after treatment with indicated doses of bemcentinib and dubermatinib for 48 h. E) Time-course proliferation curves following treatment with dubermatinib or bemcentinib in SK-LMS-1. F) Same as in D for HT-1080. *p<0.05, **p<0.01, ***p<0.001.

### 4.1.3. AXL depletion does not recapitulate the cellular effects of dubermatinib or bemcentinib

To fully validate the role of AXL in the antitumour activity of dubermatinib and bemcentinib, we generated a stable *AXL* knockdown in SK-LMS-1 and HT-1080 cells using two independent shRNA constructs (shAXL-1, shAXL-2) together with an empty-vector (EV) control, achieving a significant reduction of *AXL* mRNA and protein levels (Fig. S3). Surprisingly, AXL depletion did not reduce proliferation in either cell line (Fig. 3A), and did not modify the toxicity induced by dubermatinib or bemcentinib in SK-LMS-1 (Fig. 3B) or HT-1080 (Fig. 3C), pointing to the independence of AXL in the antitumour activity of dubermatinib and bemcentinib in these STS models and suggesting additional molecular targets may be involved.

**Fig. 3.**
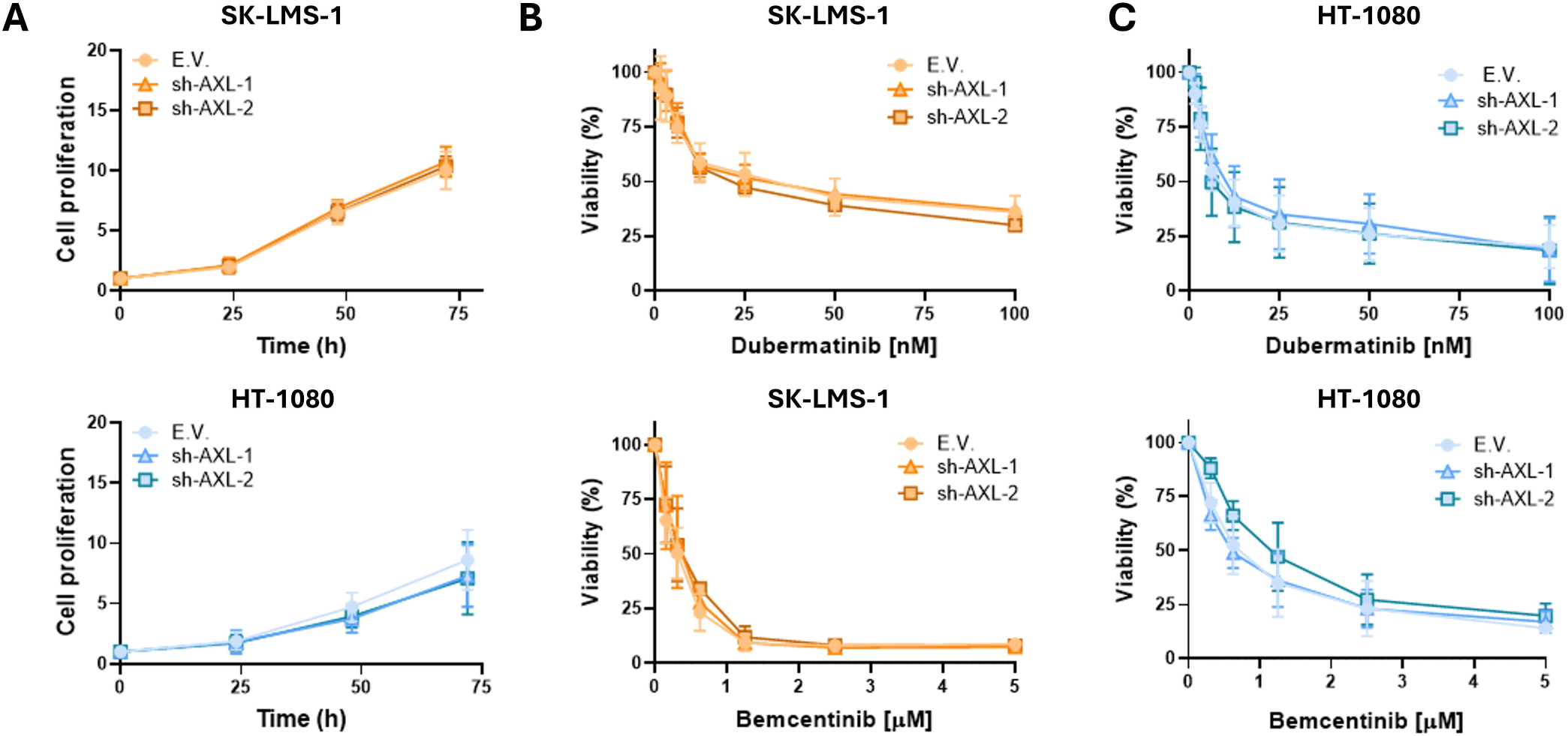
AXL knockdown does not recapitulate the effects of dubermatinib or bemcentinib. A) Proliferation curves for EV, shAXL-1 and shAXL-2 in SK-LMS-1 (top) and HT-1080 (bottom). B) Dose-response viability curves for SK-LMS-1 EV, shAXL-1 and shAXL-2 cells treated for 72 h at indicated doses of dubermatinib and bemcentinib. C) Same as in B for HT-1080.

### 4.1.4. Cell line-specific transcriptional programmes underlie AXL knockdown and drug treatment

To characterise the transcriptional changes underlying AXL depletion and pharmacological AXL inhibition, we performed RNA sequencing and analysed eight contrasts: cells treated with bemcentinib or dubermatinib and cells harbouring shAXL-1 or shAXL-2, in both HT-1080 and SK-LMS-1, each relative to its respective control (untreated cells or cells transduced with EV). Principal component analysis (Fig. 4A) showed that samples separated primarily by cell line and, within each cell line, by treatment. Spearman correlation of log2 fold changes (LFC) profiles across all contrasts clustered samples primarily by treatment rather than by cell line, although correlation values were modest throughout all comparisons (Fig. 4B), pointing to substantial differences between treatments and cell lines. *AXL* transcript levels were reduced to a comparable extent by both shRNAs in both cell lines, with no apparent compensatory effect on *TYRO3* (Fig. S4A) while MERTK was not expressed in either cell line, discarding a compensation effect within the TAM family.

**Fig. 4.**
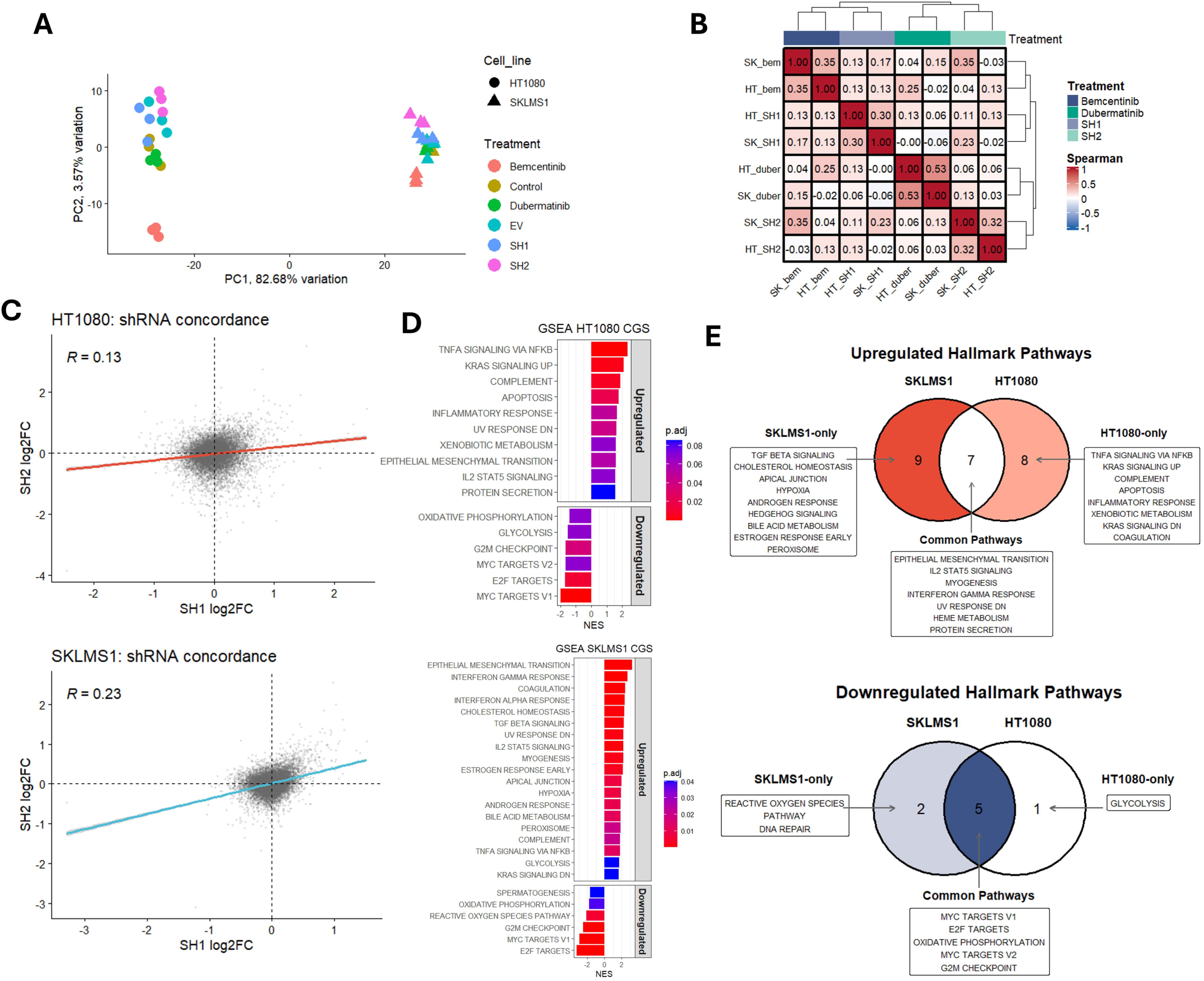
Transcriptomic characterisation of AXL knockdown and drug treatments. A) Principal component analysis of RNA-seq samples, coloured by treatment and shaped by cell line. B) Spearman correlation heatmap of LFC profiles across all contrasts. C) Concordance between shAXL-1 and shAXL-2 LFCs (Spearman correlation) in HT-1080 (top) and SK-LMS-1 (bottom). D) Hallmark GSEA of the HT-1080 (top) and SK-LMS-1 (bottom) AXL-knockdown consensus signatures. E) Venn diagrams of upregulated and downregulated Hallmark pathways shared or specific to each cell line’s consensus signature.

Because shRNA approaches are prone to off-target effects [15–17], we first compared the transcriptional changes induced by shAXL-1 and shAXL-2 within each cell line. Concordance between the two shRNAs was modest in both cell models (Spearman R = 0.13 in HT-1080 and R = 0.23 in SK-LMS-1; Fig. 4C), indicating that a substantial fraction of the transcriptional changes associated with each shRNA likely reflects off-target activity rather than on-target AXL depletion, reflected in the markedly different number of genes affected by each construct as shown in the volcano plots (Fig. S4B-C). Following common practices for mitigation of off-target effects associated with genetic perturbation methods [18,19], we derived, for each cell line, a consensus gene signature (CGS) of AXL depletion by averaging LFCs of both shRNAs for genes showing concordant direction of change. Gene set enrichment analysis (GSEA) of the resulting CGS against the MSigDB Hallmark collection identified both shared and cell line-specific pathways (Fig. 4D-E). Only 7 out of 24 upregulated Hallmark pathways were shared between both cell lines, while downregulated Hallmark pathways showed a higher relative overlap between the two lines (5 shared out of 8), yet the cross-cell-line correlation of the CGS LFC remained only partial (Spearman R = 0.25) (Fig. S4D).

### 4.1.5. Drug-induced transcriptional changes correlate weakly with the AXL-knockdown consensus signature

The number of differentially expressed genes (DEGs) varied substantially across contrasts (Fig. 5A–D and Fig. S5A), with bemcentinib inducing the broadest transcriptional response in both cell lines while dubermatinib induced a weaker response. Both drugs also induced smaller transcriptional changes in SK-LMS-1 than in HT-1080, similarly to our observations in the PCA and suggesting that SK-LMS-1 is, overall, a less transcriptionally dynamic cell line in this context. We next checked to what extent drug-induced transcriptional changes recapitulate *AXL* genetic perturbation by correlating the genome-wide LFC profile of each drug with the cell line-matched *AXL*-knockdown consensus signature. Unexpectedly, Spearman correlation was weak across all four drug-cell line combinations (bemcentinib: R = 0.208 in HT-1080 and R = 0.393 in SK-LMS-1; dubermatinib: R = 0.155 in HT-1080 and R = 0.057 in SK-LMS-1) (Fig. 5E–F). Accordingly, the consensus signature explained only a small fraction of the variance in drug-induced LFC. To discard this CGS was dominated by one of the shRNAs, we recomputed the correlation using signatures from shAXL-1 or shAXL-2 separately, which returned R² values of similar magnitude in the four drug-cell line combinations (Fig. S5B).

**Fig. 5.**
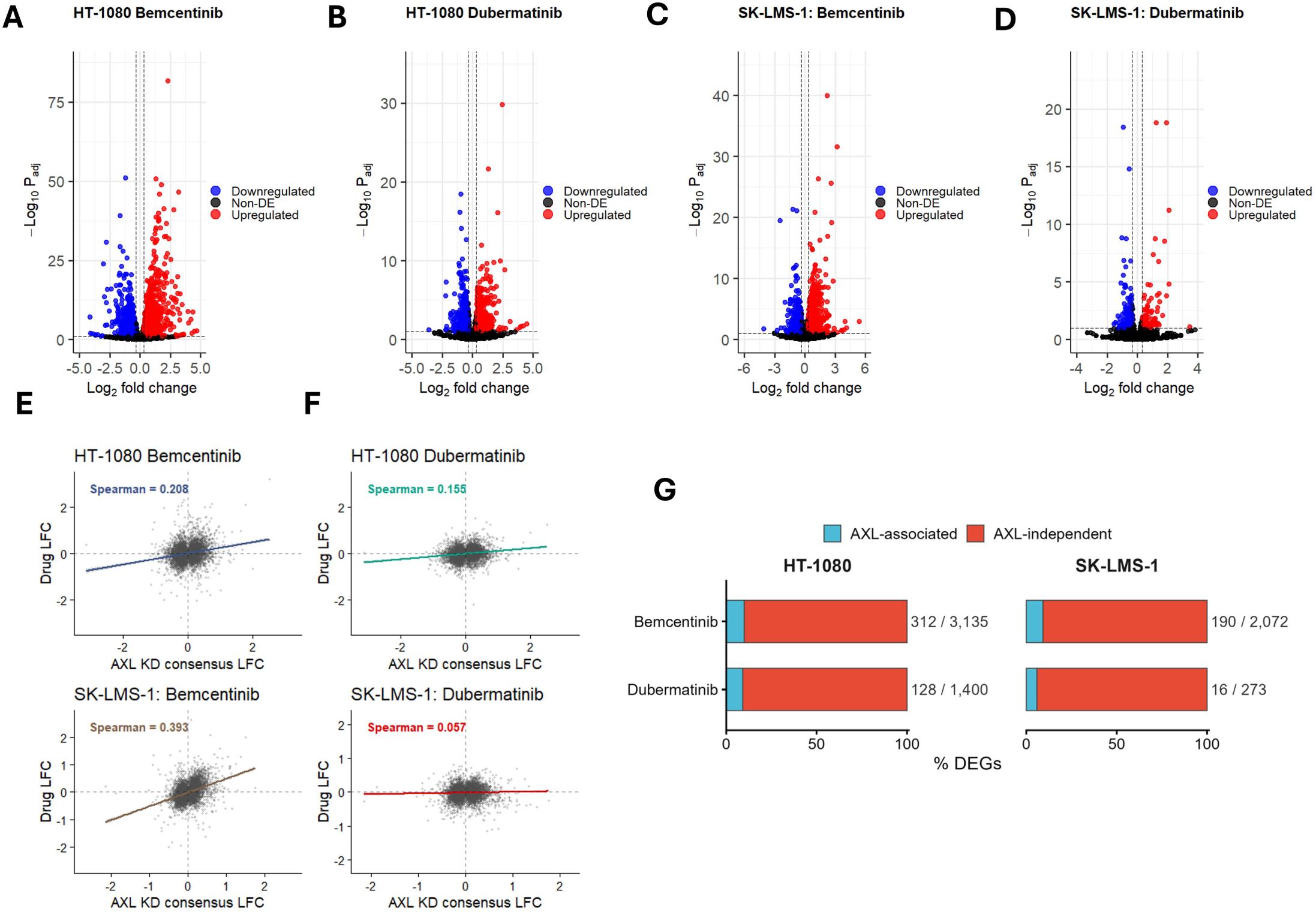
Drug-induced transcriptional changes correlate weakly with the AXL-knockdown consensus signature. A–D) Volcano plots of differentially expressed genes for bemcentinib and dubermatinib in HT-1080 (A, B) and SK-LMS-1 (C, D). E–F) Correlation between drug-induced LFC and the cell line-matched AXL-knockdown CGS LFC, for bemcentinib (E) and dubermatinib (F) in HT-1080 (top) and SK-LMS-1 (bottom). G) Number and percentage of AXL-associated versus AXL-independent DEGs for each drug-cell line combination.

Following established approaches [20], we classified drug-regulated genes as AXL-associated (concordant-direction DEGs relative to the consensus signature) or AXL-independent (all other significant DEGs) and confirmed that, in every drug-cell line combination, only a minority of DEGs could be attributed to *AXL* knockdown: 312 of 3,135 bemcentinib-regulated genes in HT-1080 (10%) and 190 of 2,072 in SK-LMS-1 (9%); 128 of 1,400 dubermatinib-regulated genes in HT-1080 (9%) and 16 of 273 in SK-LMS-1 (6%) (Fig. 5G).

### 4.1.6. AXL-independent drug effects converge on proliferation pathways in a largely cell line-specific fashion

To characterise the biological processes underlying the AXL-independent and AXL-associated components of the drug response, we performed KEGG and Gene Ontology overrepresentation analysis of each component for each drug-cell line combination, and directly compared the resulting term sets between HT-1080 and SK-LMS-1 (Fig. 6 and S6A–D). Downregulated AXL-independent genes, apart from SK-LMS-1 cells treated with dubermatinib, converged mostly on cell cycle- and DNA replication-related processes (Fig. 6B, D), with HT-1080 additionally showing enrichment for nucleotide-excision repair or homologous recombination, particularly for dubermatinib (Fig. 6D). Upregulated AXL-independent genes showed more divergent and drug- and cell line-specific profiles (Fig. 6A, C). For both drugs, the overlap of AXL-independent terms between HT-1080 and SK-LMS-1 was limited compared to the number of cell line-specific terms (Fig. 6A–D), suggesting that the biological processes mediating the AXL-independent component of the drug response are largely cell line-specific rather than a shared mechanism. In contrast, the AXL-associated gene sets showed fewer and less consistent enriched terms across the four drug-cell line combinations, potentially due to the small size of these components (Fig. S6A–D).

**Fig. 6.**
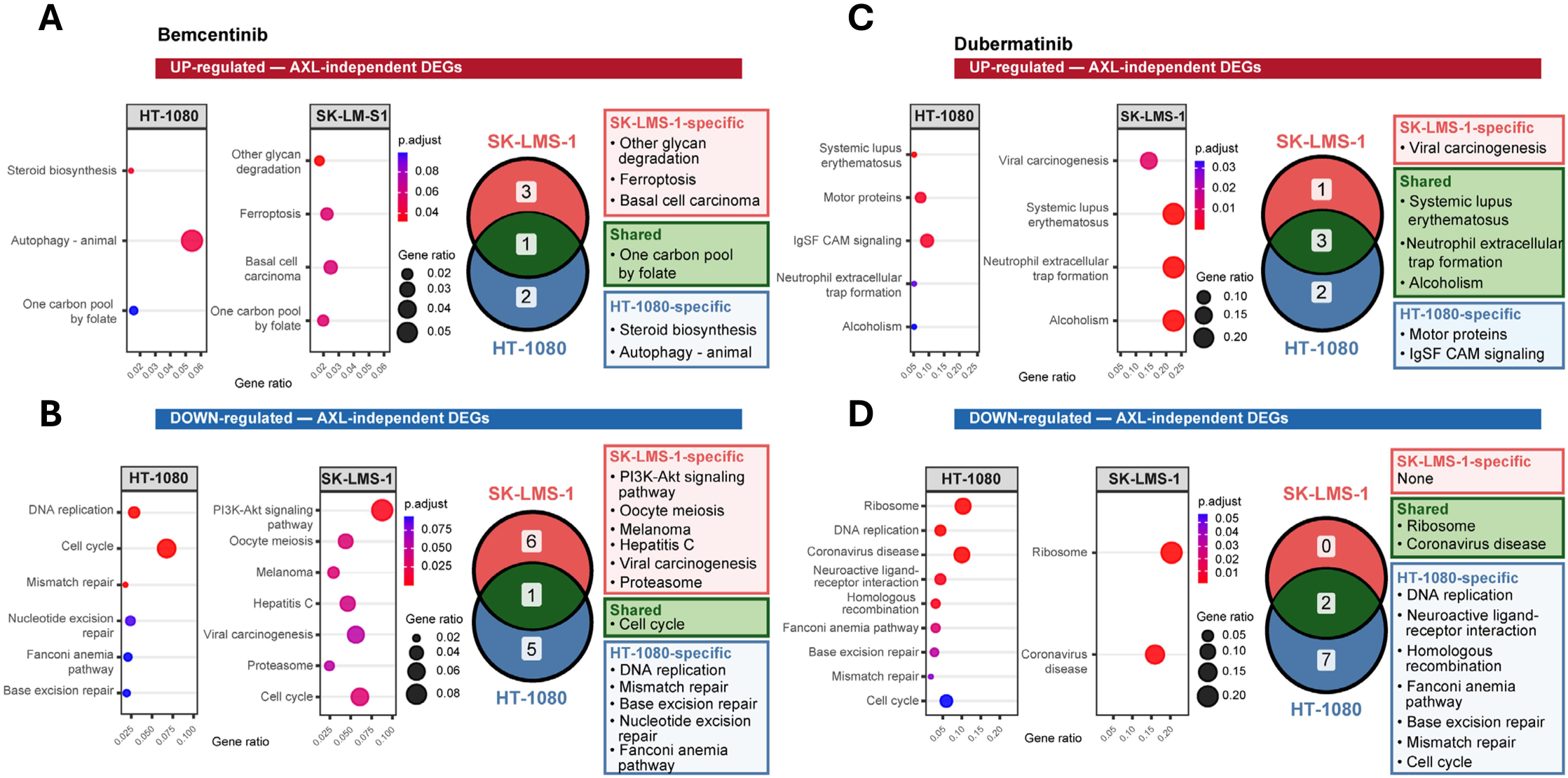
AXL-independent drug effects are largely cell line-specific. KEGG overrepresentation analysis of AXL-independent DEGs upregulated by bemcentinib (A) and dubermatinib (C), and downregulated by bemcentinib (B) and dubermatinib (D), in HT-1080 and SK-LMS-1, with Venn diagrams showing overlap of enriched terms between the two cell lines.

We functionally validated these observations by measuring cell cycle distribution (Fig. S6E). We observed that both drugs induced in HT-1080 cells an accumulation of the G2/M cell cycle phase but, surprisingly, in SK-LMS-1 only bemcentinib induced accumulation in G0/G1 phase, in contrast with HT-1080 (ANOVA interaction p value in SK-LMS-1: 0.023; in HT-1080: 0.036). These results indicate that dubermatinib and bemcentinib modulate proliferative and stress-response programmes largely independently of AXL through mechanisms that differ between drugs and STS models.

### 4.1.7. Transcriptomic responses to dubermatinib and bemcentinib resemble mitotic and PI3K–mTOR inhibition rather than AXL-inhibition

We next used decoupleR to interrogate whether the signalling pathways and transcription factors underlying the drug responses differed from those altered by AXL knockdown. Bemcentinib reduced PI3K pathway activity in both models (MLM scores −8.4 and −18.3 in HT-1080 and SK-LMS-1, respectively) whereas dubermatinib increased it (+5.9 and +3.7), with p53 activation being the only pathway shared by all six contrasts (Fig. 7A). Among the 18 transcription factors with the most variable activity across contrasts (CollecTRI), 16 had lower activity under drug treatment than in the knockdown signatures, and some even showed significant changes of opposite sign, such as NR1H3, ETV4 and EGR2 (Fig. 7B, Fig. S7A), highlighting differences in the responses triggered by each drug in each cell line.

**Fig. 7.**
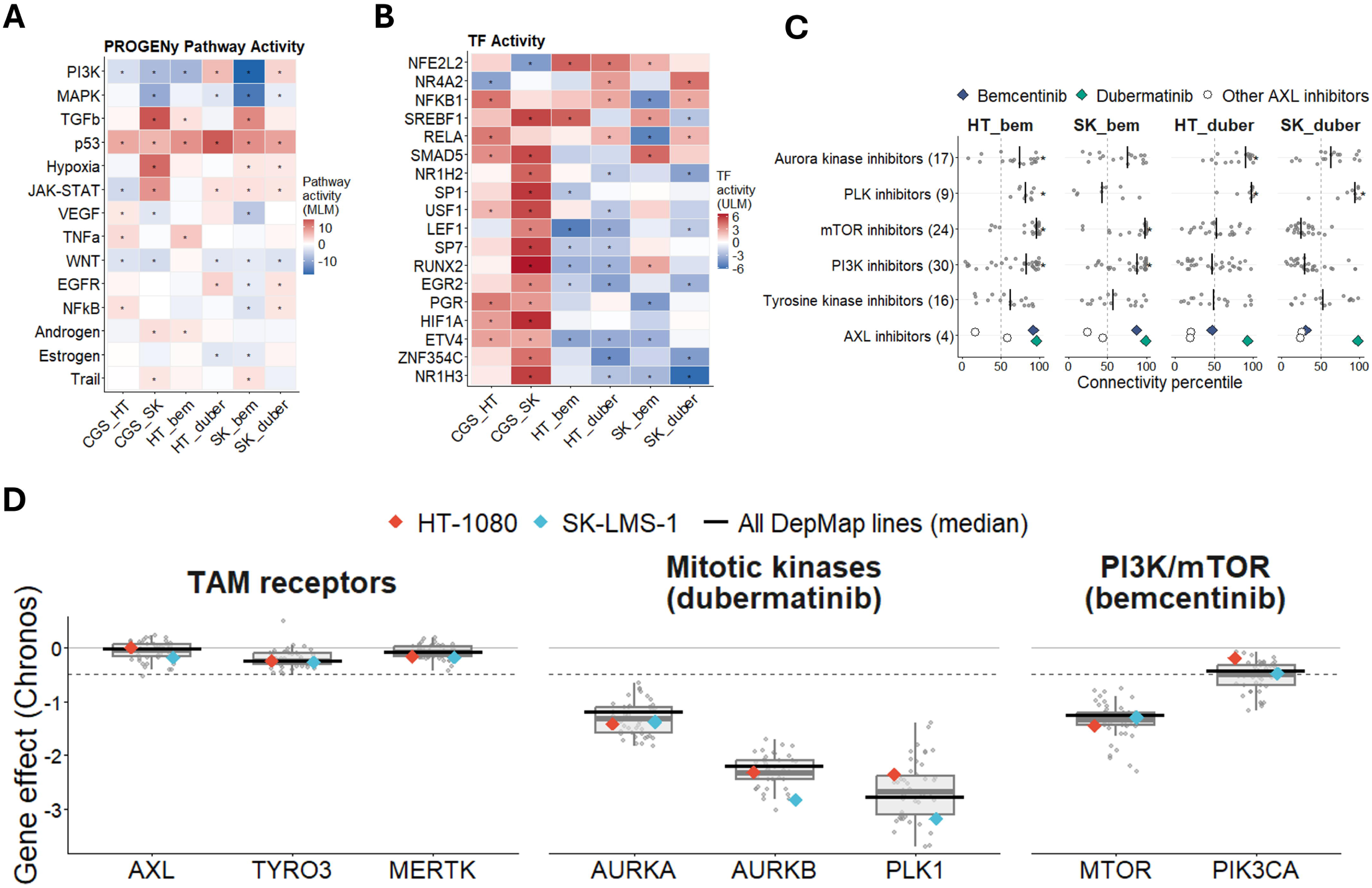
Transcriptomic responses to dubermatinib and bemcentinib resemble mitotic and PI3K–mTOR inhibition rather than AXL inhibition. A) PROGENy pathway activity inference. B) CollecTRI activity of the 18 most variable transcription factors across the AXL-knockdown consensus signatures (CGS) and drug contrasts. Asterisk means p < 0.05. C) Connectivity of each drug profile with LINCS/L1000 compounds from selected mechanism-of-action classes, expressed as connectivity percentile. Dots mean individual compounds and vertical lines mean class median, and number of compounds is shown in brackets. The tyrosine kinase inhibitor class is shown as reference, and the AXL inhibitor row shows the LINCS signatures of bemcentinib, dubermatinib and other AXL inhibitors. Asterisk means FDR < 0.05 in the class enrichment test. D) CRISPR dependency (Chronos gene effect) of TAM receptors, Aurora kinases, PLK1, MTOR and PIK3CA in 46 soft tissue sarcoma cell lines from DepMap (boxes and grey dots). Coloured symbols indicate HT-1080 and SK-LMS-1, black lines the median across all DepMap lines and the dashed line the dependency threshold (−0.5).

We then scored each drug signature against LINCS compound signatures and tested 148 mechanism of action classes for enrichment. Bemcentinib connected to mTOR and PI3K inhibitors in both cell lines while dubermatinib connected to PLK, tubulin, microtubule, CDK and cell-cycle inhibitors in both cell lines and to Aurora kinase inhibitors in HT-1080. HSP, HDAC, CHK, MEK and topoisomerase inhibitors were enriched in all four profiles, whereas the generic tyrosine kinase inhibitor class was not enriched in any contrast (FDR = 0.23–0.69) (Fig. S7B). Interestingly, the LINCS dubermatinib signature ranked above the 90th connectivity percentile for all four drug profiles, and the LINCS bemcentinib signature above the 85th percentile for both bemcentinib profiles, whereas the two other AXL inhibitors profiled in LINCS ranked very low (Fig. 7C).

In agreement with these findings, DepMap CRISPR data showed no AXL dependency in either cell line or across 46 soft tissue sarcoma lines (Fig. 7D, Fig. S7C), whereas AURKA, AURKB, PLK1 and MTOR were strong dependencies in the same models, as in most DepMap lines, and PIK3CA showed only a weak effect (Fig. 7D, Fig. S7D, E).

### 4.1.8. Neither the AXL-knockdown consensus signature nor its AXL-associated components show prognostic value in the TCGA-SARC cohort

Finally, we asked whether the transcriptional signatures are relevant from a prognostic point of view in an independent patient cohort. We scored TCGA-SARC [21] tumours by single-sample GSEA (ssGSEA) using directional (up/down) gene sets derived from the global and cell line-specific *AXL*-knockdown consensus signatures, and from the AXL-associated and AXL-independent components of each of the four drug-cell line combinations. Cox proportional-hazards models, adjusted for age and sex and corrected for multiple testing (Benjamini-Hochberg), did not identify any of the tested signatures as a significant predictor of overall survival (all FDR > 0.05) (Fig. 8A), and univariate Kaplan-Meier analysis consistently showed no significant differences between high- and low-score groups for CGS signatures nor drug-regulated AXL-independent components (all p > 0.1) (Fig. 8B–C).

**Fig. 8.**
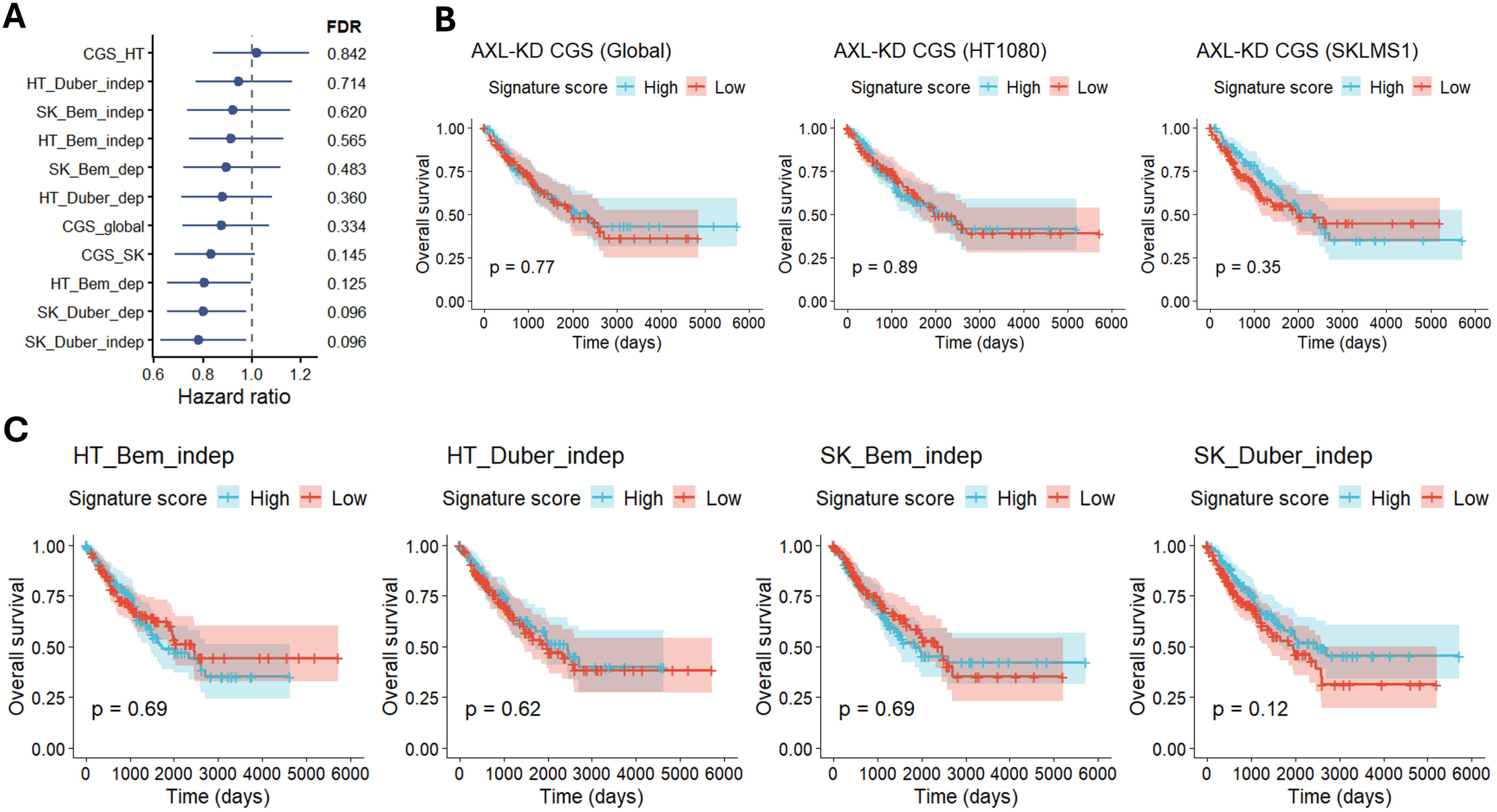
Neither the AXL-knockdown consensus signature nor its drug-associated components are prognostic in TCGA-SARC. A) Forest plot of hazard ratios (with FDR) from multivariable Cox models of overall survival for indicated ssGSEA signatures. HT, HT-1080; SK, SK-LMS-1; Bem, bemcentinib; Duber, dubermatinib; dep, AXL-associated; indep, AXL-independent. B) Kaplan-Meier curves for the global, HT-1080 and SK-LMS-1 AXL-knockdown consensus signatures. C) Kaplan-Meier curves for the AXL-independent signature of each drug-cell line combination.

The AXL-associated drug signatures showed relatively lower p-values in univariate Kaplan-Meier analysis (HT-1080 bemcentinib, p = 0.09; SK-LMS-1 dubermatinib, p = 0.11) (Fig. S8A), raising the possibility of a modest prognostic signal. However, a permutation test comparing the hazard ratio of the signature against 1,000 randomly sampled gene sets from the background of drug-regulated genes showed that the hazard ratio was not different from that expected by chance (p = 0.188; Fig. S8B), suggesting that this association is not robust to the specific gene composition of the signature.

## 5. Discussion

The identification of therapeutic targets remains a major unmet need in different tumour types including STS, and AXL has repeatedly emerged as one of the most attractive candidates across cancer types on the basis of its expression, association with drug resistance [22], and the antitumour activity of AXL inhibitors in preclinical models [3,23]. STS appeared to be a plausible subject for this strategy since AXL is highly expressed in STS relative to other TCGA tumour types, is expressed and functionally implicated in specific histological subtypes [4,5,24], and multikinase inhibitors with anti-AXL activity have shown clinical activity in sarcoma [25,26]. Our data partially confirm these observations; however, two lines of evidence in our study discard AXL as the principal driver of this activity: i) AXL knockdown did not recapitulate the antiproliferative and cytotoxic effects of dubermatinib or bemcentinib neither it modified cellular sensitivity to either drug; and ii) transcriptomic profiling showed a weak correlation between drug-induced and AXL-knockdown-induced gene expression changes, with the large majority of drug-regulated genes and pathways being AXL-independent.

These findings are in agreement with a growing body of literature questioning the target specificity of small-molecule kinase inhibitors [27,28]. Chemical proteomic profiling of clinical kinase inhibitors has repeatedly shown substantial lack of specificity at the concentrations required for their antitumour activity [29,30]. In our case, this possibility is particularly relevant for bemcentinib, which inhibits AXL with a biochemical IC50 in the nanomolar range [11], whereas the cytotoxicity we observe in STS cells requires micromolar concentrations. In this sense, it has been reported that bemcentinib induces apoptosis in cancer cells independently of AXL by blocking lysosomal acidification and recycling [31]. Interestingly, AXL-independent genes upregulated by bemcentinib were enriched in autophagy in HT-1080 and in other glycan degradation, a lysosomal pathway, in SK-LMS-1 [32] (Fig. 6A). The situation with dubermatinib is different since its activity is observed at concentrations compatible with on-target AXL inhibition, yet the lack of effect of the AXL knockdown in the toxicity of dubermatinib discards the involvement of AXL.

Unbiased connectivity analysis against LINCS/L1000 compound signatures positioned the dubermatinib response closest to mitotic inhibitor classes (PLK, tubulin and microtubule inhibitors in both cell lines and Aurora kinase inhibitors in HT-1080), in line with its reported inhibition of the Aurora kinase family [33] and with the G2/M accumulation observed in HT-1080 (Fig. S6E), the expected consequence of interfering with the Aurora A–PLK1 axis [34]. The bemcentinib response was instead closest to mTOR and PI3K inhibitors, consistent with its reduction of PI3K pathway activity. Neither compound resembled the generic tyrosine kinase inhibitor class, and DepMap data show AXL to be dispensable in both models while mTOR, PLK1 and the Aurora family are strong dependencies. We therefore propose Aurora/PLK signalling for dubermatinib and PI3K–mTOR signalling for bemcentinib as candidate mediators to be further tested experimentally. These compounds may still be of therapeutic interest in STS, but through mechanisms other than AXL inhibition which need to be better described.

On the other hand, the two shRNAs used in this work both reduced AXL substantially, yet they produced markedly divergent transcriptional profiles (Spearman R = 0.13 in HT-1080 and R = 0.23 in SK-LMS-1) and differed by more than two-fold in the number of DEGs (Fig. S4B–C, Fig. S5A). This behaviour, although striking, has been frequently reported in the literature to be associated with all kinds of genetic perturbation approaches. For instance, siRNAs directed against the same transcript have been shown to generate largely distinct signatures dominated by microRNA-like off-target regulation, driven by complementarity between the guide seed region and 3′UTRs rather than by the intended target [35,36], enough to account for a substantial fraction of the hits obtained in RNAi screens [37]. Similarly, shRNAs sharing a seed sequence resemble one another more closely than shRNAs sharing a target gene [17]. Large-scale profiling of RNAi and CRISPR perturbations has also reported that transcriptional signatures obtained with different molecules against the same gene correlate poorly despite comparable on-target silencing, and averaging constructs into a consensus signature is required to improve the precision of the on-target component [17,38]. A similar scenario has been observed for CRISPRi, where guides directed at identical targets yield non-overlapping sets of differentially expressed genes and even different cellular phenotypes, attributable to off-target activity [39]. Our consensus signature strategy was adopted precisely to mitigate this limitation.

Another unexpected observation in our data was the degree of cell line specificity in both the AXL-knockdown and drug-induced transcriptional responses. HT-1080 consistently showed a broader transcriptomic response to both shRNA-mediated knockdown and pharmacological treatment than SK-LMS-1. Additionally, both drugs down-regulated cell cycle pathways and increased the G2/M fraction in HT-1080 cells, whereas in SK-LMS-1 cell cycle pathways were down-regulated only by bemcentinib, which induced accumulation in G0/G1 instead (Fig. 6B-D, Fig. S6E), pointing to different molecular mechanisms in the two cell lines. This discrepancy may reflect differences in transcriptional dynamics, chromatin state, or compensatory signalling between the two lines, and cautions against generalising AXL-dependence conclusions from a single STS model, which in fact reflects the high heterogeneity of this group of tumours [21]. In the same line, the limited overlap in AXL-independent pathways between the two cell lines (Fig. 6) indicates that off-target and AXL-independent mechanisms of action of drugs are substantially cell type-dependent, with direct implications for identifying which STS subtypes might benefit from these therapies. Additionally, the absence of prognostic value of AXL in the TCGA-SARC cohort, in contrast with observations reported in other tumour types and even in certain sarcoma subtypes [4,5], suggests the role of AXL as prognostic marker may also be carefully revisited.

In summary, our data support a potential antitumour role for dubermatinib and bemcentinib in STS, but indicate that this therapeutic activity is largely independent from AXL inhibition. On the whole, these findings underscore the need for caution in interpreting the antitumour effects of AXL-targeted kinase inhibitors, and point to mitotic and PI3K–mTOR signalling pathways as candidate mediators whose experimental validation should be a priority to exploit their therapeutic potential in STS.

## 6. Methods

### 6.1. In silico analysis of TCGA and CCLE datasets

Gene expression, copy-number, mutation and structural-variant data across pan-cancer TCGA cohorts were obtained from the Genomic Data Commons (GDC)[40] via the TCGAbiolinks R package [41]. AXL expression was normalised to counts per million (CPM) using per-sample library size and transformed to log2(CPM + 1).

Survival analyses in TCGA-SARC were performed using clinical data from GDC and patients were dichotomised into high- and low-AXL groups by the median log2(CPM + 1) value. Kaplan-Meier curve was generated with survival [42] and survminer [43] R packages, and hazard ratios were estimated by multivariable Cox proportional-hazards models adjusting for age and sex.

AXL mRNA expression across tumour-derived cell lines was obtained from the Cancer Cell Line Encyclopedia (CCLE/DepMap) [44].

### 6.2. Cell lines and reagents

HT-1080 (RRID: CVCL_0317; fibrosarcoma), SK-LMS-1 (RRID: CVCL_0628; leiomyosarcoma) and HEK293T (RRID: CVCL_0063) human cell lines were obtained from ATCC (LGC Standards S.L.U., Barcelona, Spain) and cultured in Dulbecco’s modified Eagle’s medium (DMEM) supplemented with 10% fetal bovine serum and 1% glutamine plus antibiotics, at 37°C in a humidified atmosphere with 5% CO₂.

Dubermatinib (TP-0903) and bemcentinib (R428) were obtained from MedChemExpress, (EURODIAGNOSTICO S.L., Madrid, Spain), and dissolved in DMSO, aliquoted and stored at - 80°C. Puromycin (Merck) was dissolved in double-distilled water, aliquoted, and stored at -20°C.

### 6.3. shRNA knockdown

AXL knockdown was generated in HT-1080 and SK-LMS-1 cells using two independent short-hairpin RNA constructs targeting AXL: Human pLKO.1-puro-shRNAAXL (shAXL-1: TRCN0000001039 and shAXL-2: TRCN0000001040) and an empty-vector (EV) control, delivered by lentiviral transduction and selected with 1 μg·mL−1 puromycin for 72 h. Knockdown efficiency was confirmed at both protein level by western blot and mRNA level by RT-qPCR.

### 6.4. Cell viability and proliferation measurements

Cell viability following dubermatinib or bemcentinib treatment was assessed by MTT assay seeding 10^4^ cells/well in 24-well plates and treated 24 h later. Toxicity was measured after 72 h of treatment. For cell proliferation assays, 10^4^ cells/well were seeded in 24-well plates and proliferation was analysed 1, 2 and 3 days later by an MTT-based assay as previously described [45]. Data shown are the average of three independent experiments performed in triplicate cultures.

For cell cycle analysis, 10^5^ cells were seeded in 6 cm plates 24 h prior to treatment and cell cycle was analysed 24 h after treatment as previously described [46]. For apoptosis induction, 10^5^ cells were seeded in 6 cm plates, 24 h later cells were treated with drugs and, after 48 h, apoptosis was measured with Annexin V-FITC (Immunostep) following the manufacturer’s instructions. Samples and data were processed in a MACSQuant Analyzer 10 (Miltenyi Biotec). Data shown are the average of three independent experiments.

Statistical comparisons used two-way ANOVA followed by Bonferroni’s multiple comparison t-test, with significance thresholds indicated in figure legends (*p < 0.05, **p < 0.01, ***p < 0.001).

### 6.5. RNA extraction, RT-qPCR and mRNA sequencing

Total RNA was extracted from HT-1080 and SK-LMS-1 cells treated with dubermatinib, bemcentinib or vehicle control (DMSO), and from AXL-knockdown (shAXL-1, shAXL-2) and EV-control for each cell line using NZY Total RNA Isolation kit. Primers for all target sequences were designed by using the NCBI Primer-BLAST tool and purchased from Merck as DNA oligos. Primer sequences were AXL-Forward: GACATAGGGCTAAGGCAAGAGG, AXL-Reverse: CGAGAAGGCAGGAGTTGAAGG, GAPDH-Forward: TCGTGGAAGGACTCATGACCA, GAPDH-Reverse: CAGTCTTCTGGGTGGCAGTGA. Data shown are the average of, at least, three independent experiments performed in triplicate. mRNA sequencing library preparation and sequencing (20 million PE150 reads per sample) were performed by Novogene.

### 6.6. Western blotting

Cells were collected in RIPA lysis buffer. Protease and phosphatase inhibitors (Merck) were added prior to lysis. Protein quantification and western blotting was performed as previously described [47]. Fifty micrograms of protein was loaded onto appropriate percentage SDS/PAGE, transferred to PVDF membranes using semidry Pierce Power Blot (Thermo Fisher Scientific, Alcobendas, Madrid, Spain) and blotted against different proteins via specific antibodies. Antibodies were detected by enhanced chemiluminescence (Amersham Thermo Fisher Scientific) in a LAS-3000 system (FujiFilm, Alcobendas, Madrid, Spain). Antibodies used for western blot were AXL (#8661 Cell Signaling Technology), vinculin (#V9264 Sigma-Aldrich), Anti-mouse-HRP (#7076 Cell Signaling Technology) and Anti-rabbit-HRP (#7074 Cell Signaling Technology).

### 6.7. Differential gene expression analysis

Raw FASTQ files were aligned to the human reference genome GRCh38 with STAR [48] using standard parameters and the resulting BAM files were sorted and indexed with SAMtools [49] in the CSIC High-Performance Computing service. Gene-level read counts were then quantified with featureCounts (RSubread) [50] in R and analysed using DESeq2 [51]. Genes with less than 30 counts in all replicates of at least one experimental group were discarded. A Wald test with Benjamini-Hochberg correction was used to evaluate significance and genes were considered differentially expressed at padj < 0.1 and |LFC| > 0.33 for our exploratory analysis. Gene symbols and Entrez identifiers were annotated using the org.Hs.eg.db R package [52].

### 6.8. Consensus AXL-knockdown gene signature (CGS)

For each cell line, a consensus gene signature (CGS) of AXL depletion was derived by averaging log2 fold-change values across the two independent shRNAs relative to the EV control, restricted to genes showing concordant direction of change. A global consensus signature was similarly derived by averaging log2 fold-changes across all four knockdown contrasts (two shRNAs in each cell line), again retaining genes with a consistent direction of change. Signature concordance was quantified using Spearman’s rank correlation coefficient and the coefficient of determination (R²) from linear regression of drug LFC and CGS LFC. 95% confidence intervals for R² were obtained by resampling of genes (1,000 iterations) [53].

### 6.9. Gene set enrichment and overrepresentation analyses

Gene set enrichment analysis (GSEA) against the MSigDB Hallmark collection was performed using GSEA function from clusterProfiler [54]. Overrepresentation analysis of Gene Ontology (Biological Process) and KEGG pathway terms was performed using clusterProfiler’s enrichGO and enrichKEGG functions, with the genes detected in the contrasts used as the statistical background.

#### Transcription factor and pathway activity inference

Regulatory activities were inferred with decoupleR [55] from a matrix of log2 fold changes from the eight experimental contrasts and the two cell line-specific consensus signatures. Transcription factor activities were estimated with a univariate linear model against the CollecTRI collection [55], retrieved from OmniPath [56]. Pathway activities were inferred with a multivariate linear model against the PROGENy top-500 footprint models [57]. Significant transcription factors (p < 0.05) in at least two contrasts were ranked by the standard deviation of their activity across the consensus signatures and drug contrasts.

#### Connectivity analysis against LINCS compounds

Level-5 signatures of the LINCS 2020 release [58] were obtained for compound perturbations and restricted to the 978 landmark genes. Each query was the log2 fold-change profile of the corresponding contrast, mapped to Entrez identifiers and intersected with the landmark space, and connectivity was computed as its Spearman rank correlation with each LINCS signature. Compound-level scores were summarised as the mean correlation across the signatures annotated to each compound and ranked. Mechanism-of-action classes with at least five compounds were tested by Wilcoxon rank-sum test (with Benjamini-Hochberg correction) against the remaining compounds.

#### CRISPR dependency analysis

Gene effect (Chronos) scores from the DepMap CRISPR screen [59,60] were retrieved for 1,196 cell lines, of which 46 were annotated to the soft tissue lineage and dependency was defined by the conventional threshold of Chronos < −0.5.

### 6.10. ssGSEA signature scoring and survival analysis in TCGA-SARC

Directional (upregulated/downregulated) gene signatures were derived for the global and cell line-specific AXL-knockdown consensus profiles, and for the AXL-associated and AXL-independent components of each of the four drug-cell line combinations. Single-sample GSEA (ssGSEA) scores for each signature were computed on normalised TCGA-SARC expression data using ssgseaParam function from the GSVA R package [61] and a directional score (up minus down) was derived for each sample and signature. Cox proportional-hazards models regressing overall survival on the directional score, adjusted for age and sex, were fit for each signature, with Benjamini-Hochberg correction for multiple testing across signatures. A permutation test (1,000 permutations, using random gene sets of matched size from the background of genes detected in the relevant contrasts) was used to evaluate the significance and robustness of the relevant signatures.

## 8. Author contributions

DMFA: designed the study, performed all computational analyses and wrote the original manuscript; CGG and PFA: performed and analysed the *in vitro* experiments, contributed to manuscript revision and data visualization. VJSAL: provided support for computational analyses and manuscript writing and reviewing; RSP: Obtained funding, contributed to original manuscript design, and provided support during manuscript editing; MJRH, AFA, BB, FJC and NMG provided support during project and manuscript editing. All authors have reviewed and approved the final manuscript.

