## Supplementary figures and images for "Revisiting AXL as a therapeutic target in soft tissue sarcoma: transcriptomic profiling reveals largely AXL-independent activity of dubermatinib and bemcentinib"

### Supplementary Figure 1

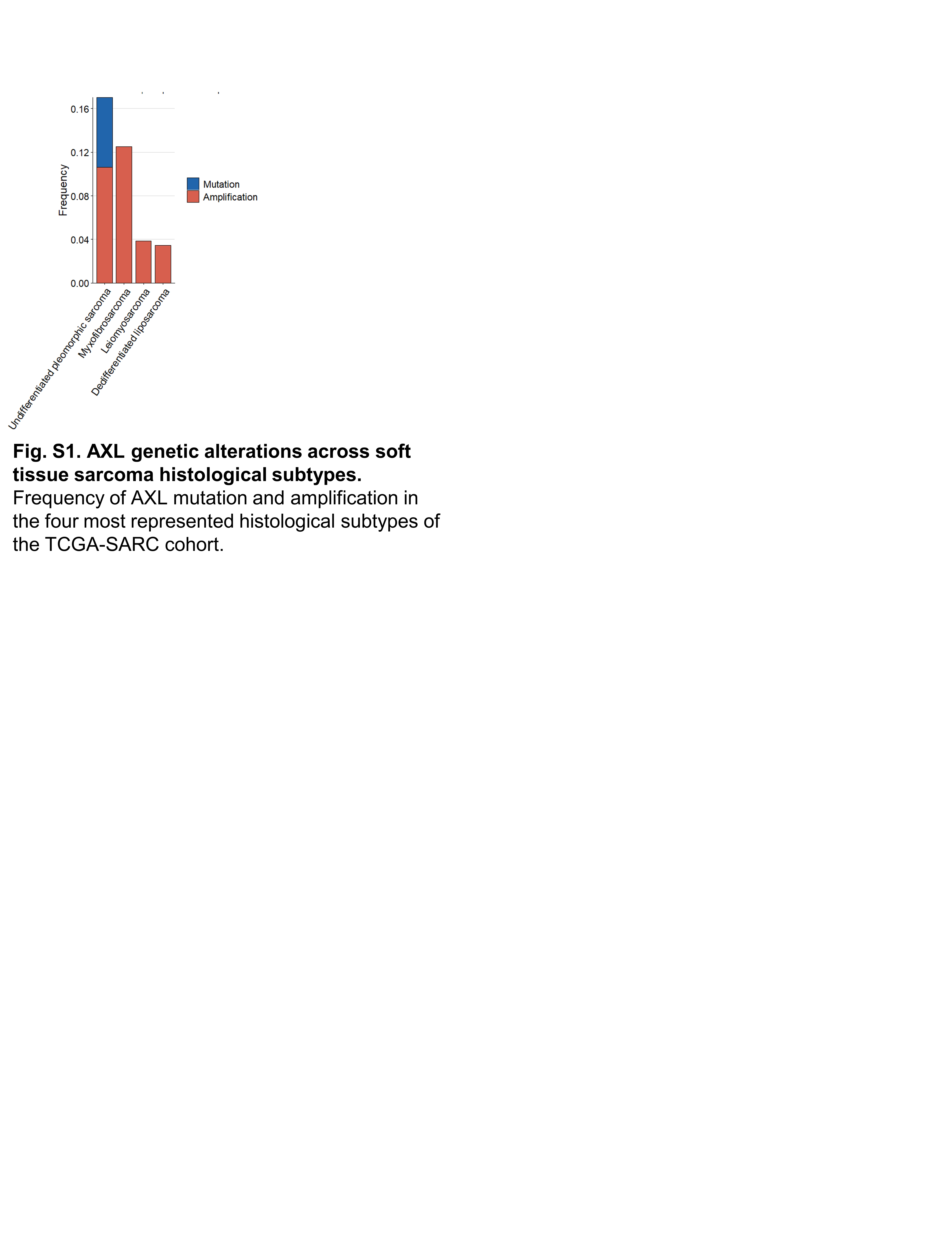

### Supplementary Figure 2

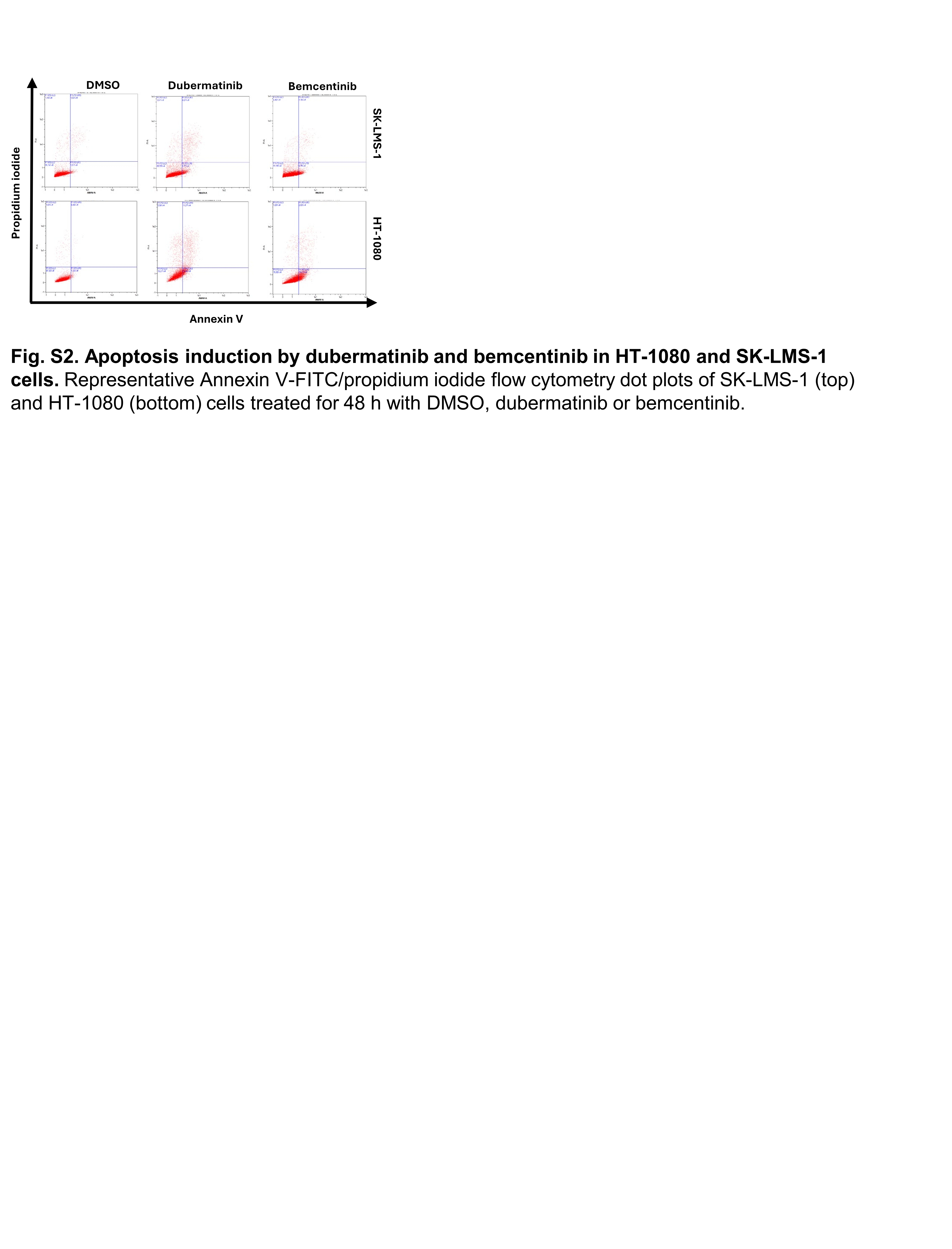

### Supplementary Figure 3

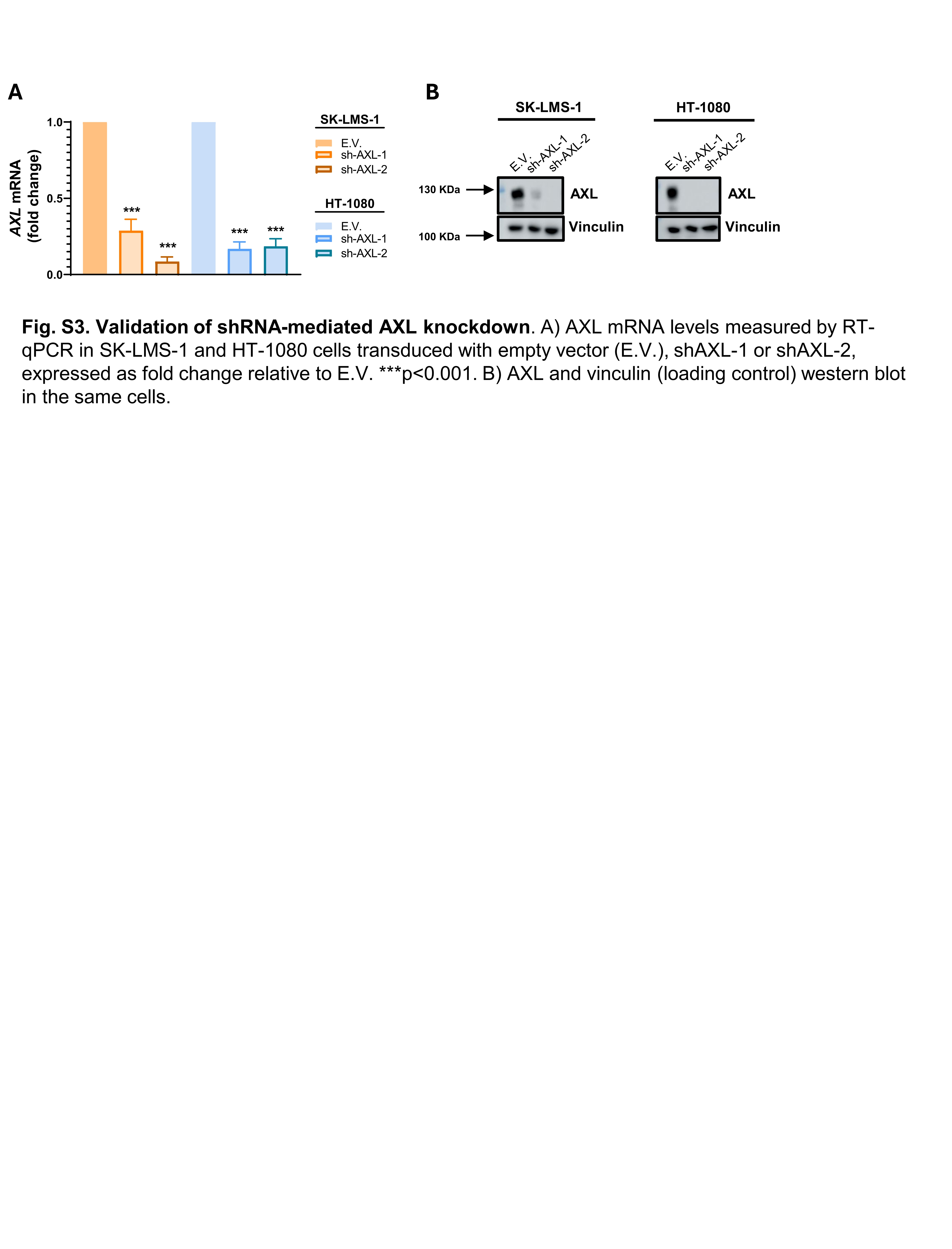

### Supplementary Figure 4

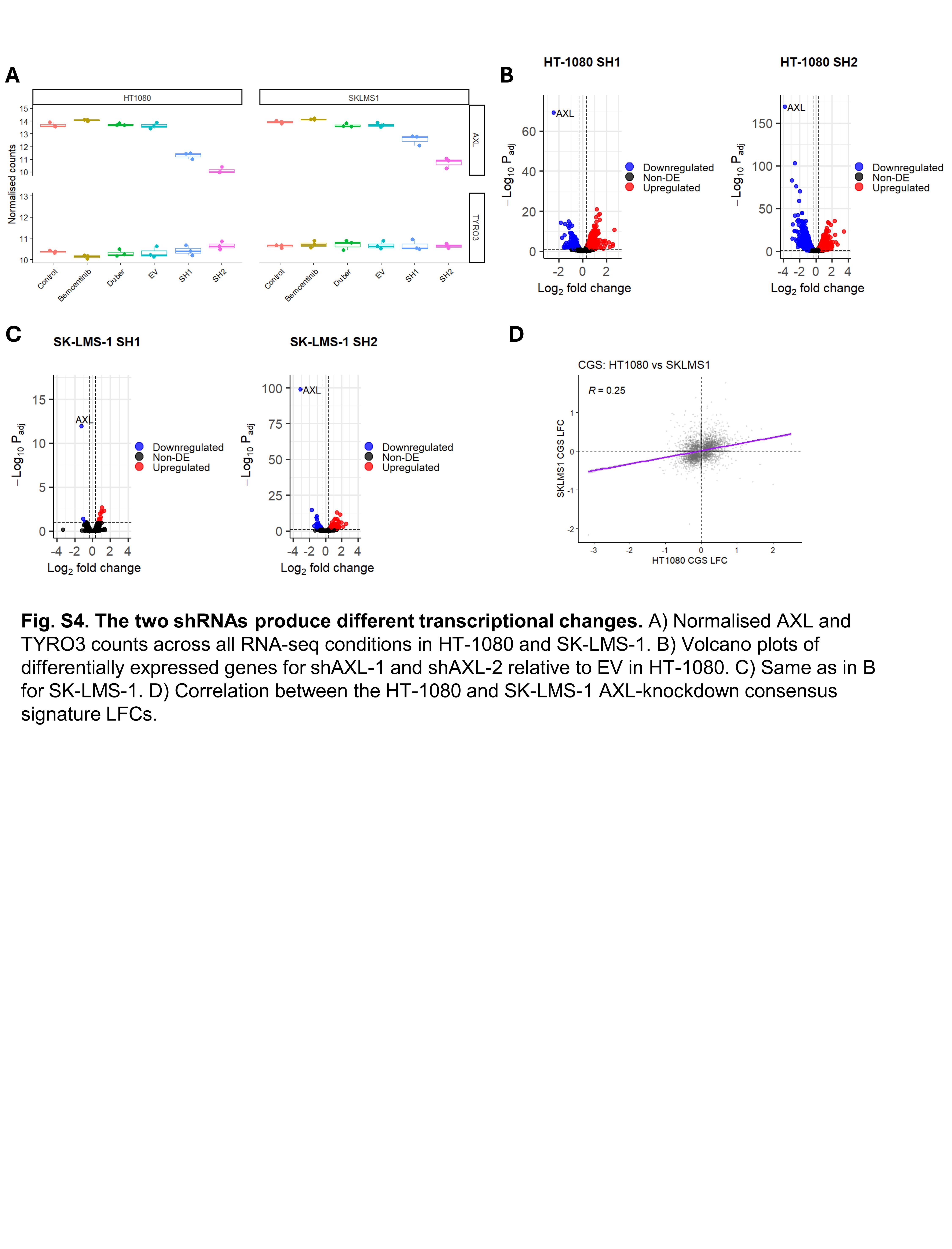

### Supplementary Figure 5

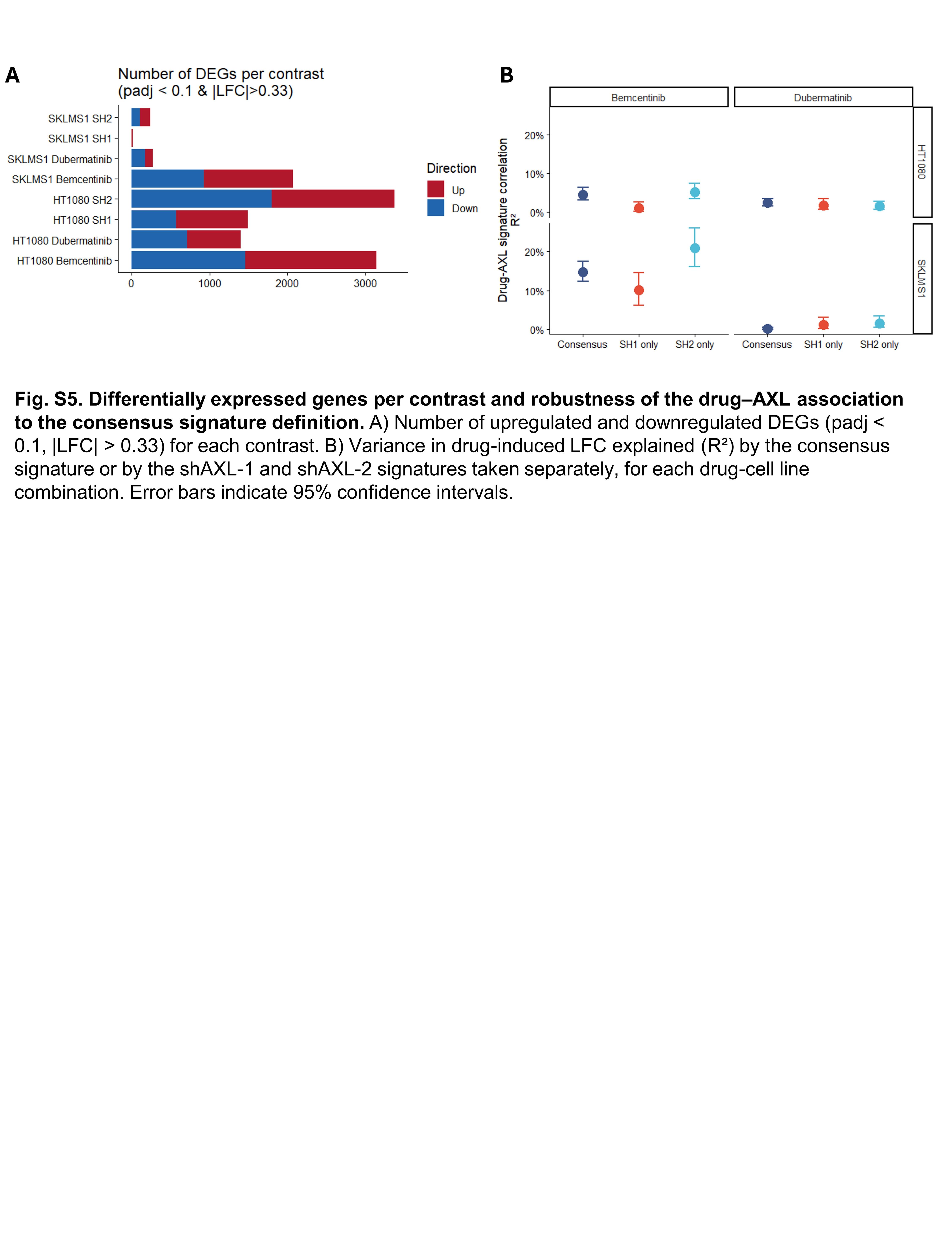

### Supplementary Figure 6

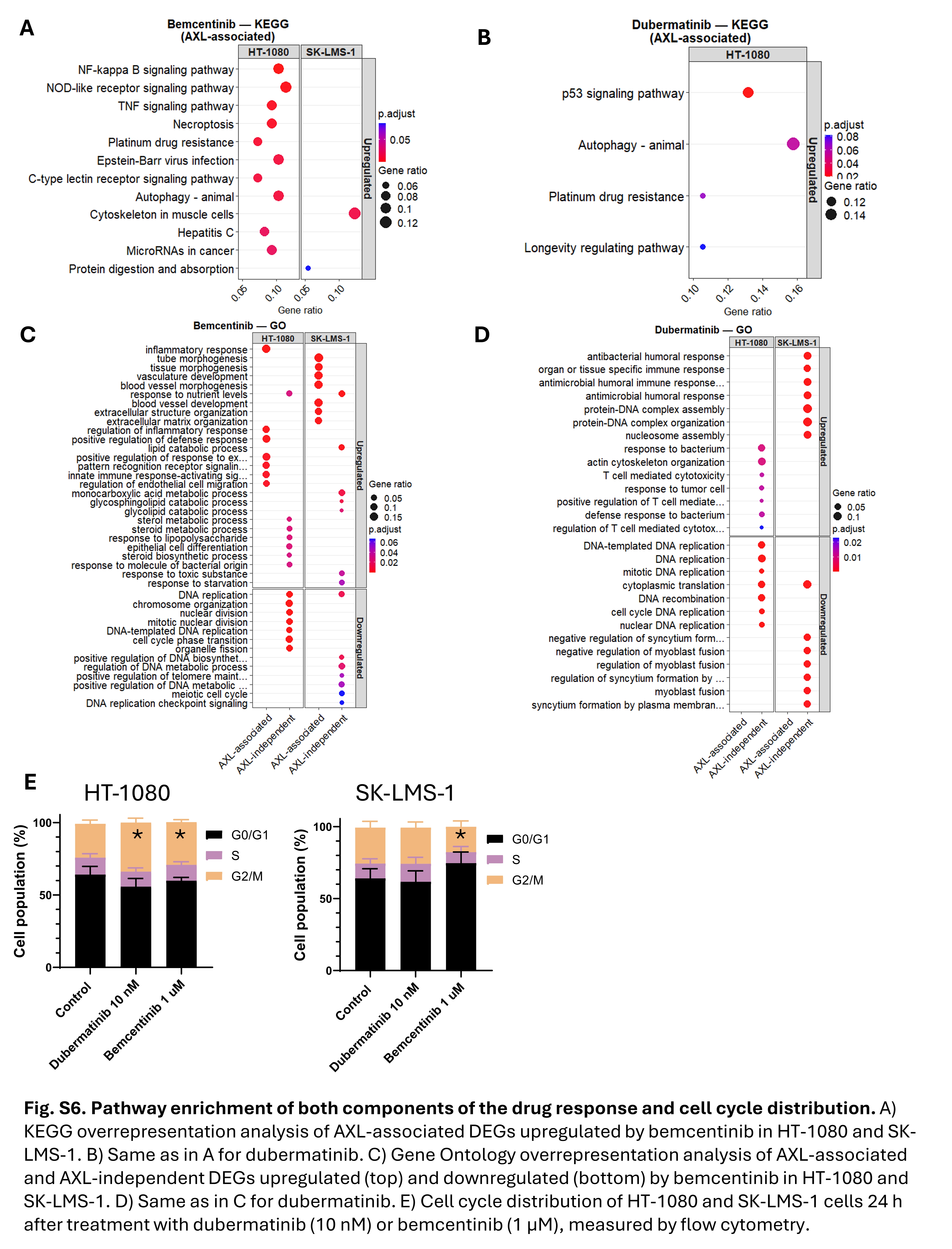

### Supplementary Figure 7

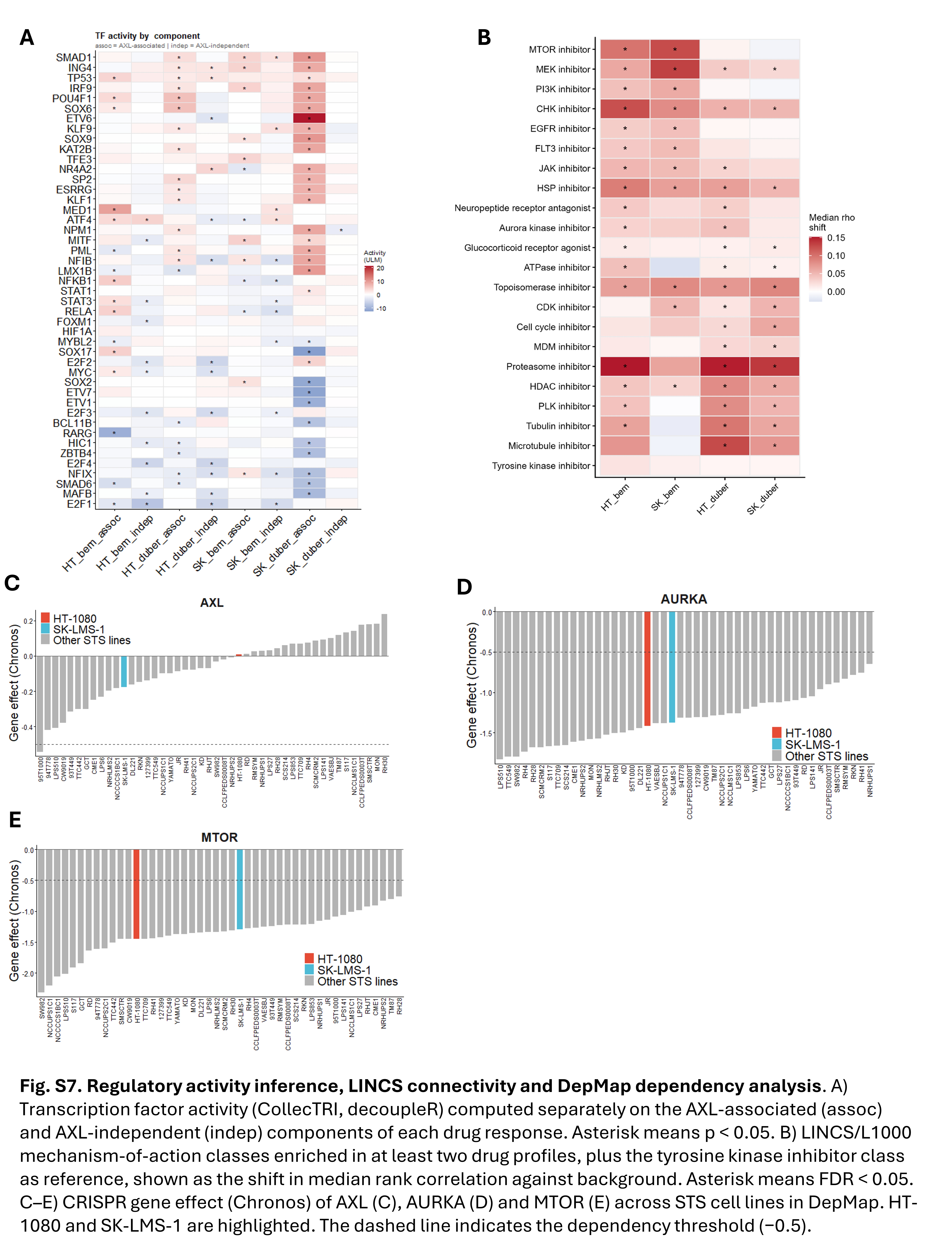

### Supplementary Figure 8

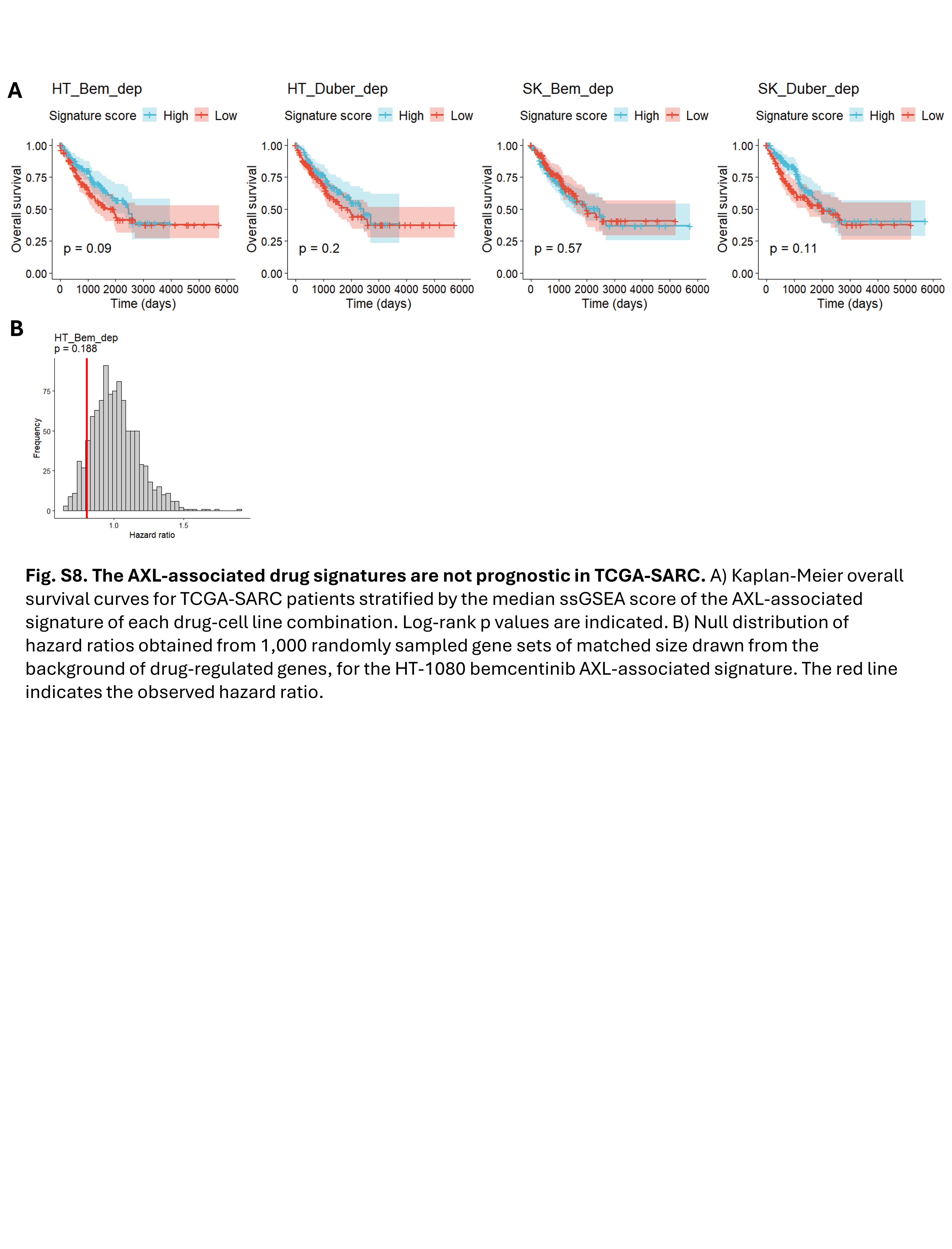
